# Visual system hyperexcitability and compromised V1 receptive field properties in early-stage retinitis pigmentosa in mice

**DOI:** 10.1101/2022.05.16.492079

**Authors:** Henri Leinonen, David C Lyon, Krzysztof Palczewski, Andrzej T Foik

## Abstract

Inherited retinal degenerative diseases are a prominent cause of blindness. Even though mutations causing death of photoreceptors are mostly known, the pathophysiology downstream in the inner retina and along the visual pathway is incompletely characterized in the earliest disease stages. Here we investigated retinal, midbrain and cortical visual function using electroretinography (ERG), the optomotor response (OMR), visual evoked potentials (VEPs), respectively, and single unit electrophysiology at the primary visual cortex (V1) in light-adapted juvenile (∼ 1-month-old) and young adult (3-month-old) *Rho*^P23H/WT^ mice, representative of early-stage retinitis pigmentosa (RP). Photopic ERG revealed up to ∼ 30 % hypersensitivity to light in *Rho*^P23H/WT^ mice, as measured by the light intensity required to generate half-maximal b-wave (I_50_ parameter). *Rho*^P23H/WT^ mice also showed increased optomotor responses towards low spatial frequency drifting gratings, indicative of visual overexcitation at the midbrain level. At the V1 level, VEPs and single-cell recordings revealed prominent hyperexcitability in the juvenile *Rho*^P23H/WT^ mice. Mean VEP amplitudes for light ON stimuli were nearly doubled in 1-month-old *Rho*^P23H/WT^ mice compared to controls, and more than doubled for light OFF. Single-cell recordings showed a significantly increased spontaneous V1 neuron firing in the *Rho*^P23H/WT^ mice, and persistent contrast and temporal sensitivities. In contrast, direction selectivity was severely compromised. Our data suggest that during early RP, the visual pathway becomes hyperexcited. This could have both compensatory and deleterious consequences for visual behavior. Further studies on the mechanisms of hyperexcitability are warranted as this could lead to therapeutic interventions for RP.

**Significance statement:** Lost retinal function in many blinding retinal degenerative disorders could soon be alleviated by advanced therapies that restore photoreception. However, it is unknown whether a visual system rewired downstream of the photoreceptors can process signals adequately. We studied the functional consequences of early rod death along the visual pathway in young retinitis pigmentosa (RP) mice. Photopic inner retina responses were moderately hypersensitized in the electroretinograms of RP mice. Reflex-based visual behavior and visual cortex electrophysiology showed hyperexcitability. Some aspects of complex visual processing were remarkably resistant to degeneration, whereas others were severely impacted. We conclude that the visual system adapts to lost photoreception by increasing sensitivity, but simultaneously becomes detrimentally hyperexcited. Mechanistic understanding could lead to therapeutic preservation and restoration of vision.

## Introduction

Consequences of total or near total loss of outer retinal photoreceptors, rods and cones, to the inner retina function have been widely studied using animal models of retinal degenerative diseases (D’Orazi *et al*., 2014; Strettoi, 2015; Pfeiffer *et al*., 2020; Strettoi *et al*. 2022, Int J Mol Sci). With respect to clinical correlates, corresponding anatomical data have been obtained from patient retinas *post mortem* (Jones *et al*., 2016). Such studies have demonstrated robust rewiring of neural connections, commonly referred to as ” remodeling”, leading to dramatically increased spontaneous neural activity, decreased signal-to-noise ratios, and attenuated light responses in the inner retina (Toychiev *et al*., 2013; Trenholm and Awatramani, 2015; Haselier et al., 2017; Telias *et al*., 2020). Much concern has been raised that retinal remodeling may preclude restoration of visual function even if augmentation of photoreception by various advanced therapies could be achieved (Marc *et al*., 2014; Reh, 2016; Foik *et al*., 2018; Suh *et al*., 2020). By default, this refractoriness would be particularly problematic in the adult organism with limited capability for neural plasticity and presumed inability to adapt to and properly process the restored sensory signals.

Much less attention has been directed to investigating functional outcomes of early retinal degeneration and milder disease stages, even though most patients with retinal degeneration suffer from a partial loss of vision, and some may never become legally blind (Berson *et al*., 2002; Xu *et al*., 2020). Studies focusing on early disease stages could help us understand the triggers and mechanisms of retinal remodeling and lead to increased ability to develop vision restoration interventions. Equally important, increased mechanistic understanding could help us design therapies to increase quality of life in patients with partial loss of vision due to retinal degeneration.

Most inherited retinal degenerations are classified under the umbrella term retinitis pigmentosa (RP). In a typical RP case, primarily rod photoreceptors are affected first; consequently the disease first manifests as loss of night vision and shrinking of the visual field (Hartong *et al*., 2006). Central and color vision impairment manifest only at a much more advanced disease state. RP phenotype has been studied extensively with respect to the primary insult, rod degeneration and dysfunction. Investigation of the cone-associated phenotype has gained somewhat less attention, even though the secondary cone degeneration may be a more realistic target for conventional pharmacological interventions. In fact, preservation of cone-pathway function could retain the most important attributes of vision for humans, such as color vision and high visual acuity. Another gap in knowledge exists in how visual signals are modulated in the brain during early RP. This is crucial as the complex circuitry in the inner retina, which ultimately sends the visual signals to the rest of the brain, rewires early in RP (Soto and Kerschensteiner, 2015). In the current study, we addressed these issues by using an established animal model of autosomal dominant RP (heterozygote *Rho*^P23H/WT^ mice) before cone degeneration, and by recording photopic electroretinography, behavioral optomotor responses, and primary visual cortex responses to light ON-OFF and multimodal pattern stimuli.

## Materials and Methods

### Animals

A mouse model of autosomal dominant retinitis pigmentosa (RP) was used in this study. The mouse carries a P23H mutation in the rhodopsin gene, causing a phenotype of extremely fast rod photoreceptor degeneration in homozygous mutants (*Rho*^*P23H/P23H*^), and an intermediately fast progressing rod degeneration in heterozygous mutants (*Rho*^*P23H/WT*^) (Sakami *et al*., 2011, 2014). This report mainly focuses on the heterozygous mutants. As is common for the RP phenotype (Hartong *et al*., 2006), cone photoreceptor degeneration is much delayed in *Rho*^*P23H/WT*^ mice (Sakami et al., 2011, 2014). The *Rho*^*P23H/WT*^ mice were generated by crossbreeding *Rho*^*P23H/P23H*^ mice with wild-type (WT) C57BL/6J mice (The Jackson Laboratory, stock # 000664). Age-matched WT mice were used as controls. Both male and female mice were used that were group-housed in a standard vivarium using *ad libitum* feeding. The light-dark cycle was set at 12 hr / 12 hr (lights on 6:30 am, lights off 6:30 pm). Cone-transducin knockout mice (*Gnat2*^*-/-*^, a kind gift of Dr. Marie Burns, UC Davis), that lack cone-mediated function but do not show retinal degeneration (Ronning *et al*., 2018), were used for calibration of photopic ERG background (Supplementary Figure 1). In all procedures, animal subjects were treated following the NIH guidelines for the care and use of laboratory animals, and the ARVO Statement for the Use of Animals in Ophthalmic and Vision Research; and under a protocol approved by the Institutional Animal Care and Use Committee of UC Irvine (AUP-18-124).

### Electroretinography

The electroretinography (ERG) was performed under standard laboratory lighting conditions using a Diagnosys Celeris rodent ERG device (Diagnosys, Lowell, MA), with some modifications from a previous protocol (Orban *et al*., 2018). The mice were anesthetized with ketamine (100 mg/kg, KetaVed; Bioniche Teoranta, Inverin Co., Galway, Ireland) and xylazine (10 mg/mg, Rompun; Bayer, Shawnee Mission, KS) by intraperitoneal injection, and their pupils were dilated with 1% tropicamide (Tropicamide Ophthalmic Solution USP 1%; Akorn, Lake Forest, IL), and thereafter kept moist with 0.3% hypromellose gel (GenTeal; Alcon, Fort Worth, TX). Light stimulation was produced by an in-house scripted simulation series in Espion software (version 6; Diagnosys). The eyes were stimulated with a green light-emitting diode (LED) (peak 544 nm, bandwidth 160 nm) or with a UV LED (peak emission 370 nm, bandwidth 50 nm). While green stimulation was used, the steady rod-suppressing background light consisted of 200 cd/m^2^ red (peak 630 nm, bandwidth 100 nm) and 100 cd/m^2^ UV. The background during UV stimulation consisted of 200 cd/m^2^ red and 100 cd/m^2^ green. Steady red light at 200 cd/m^2^ suppresses the ERG signal practically fully in *Gnat2*^*-/-*^ mice (Supplementary Figure 1). However, a steady strong UV or green background at 100 cd/m^2^ was further added to facilitate rod suppression during monochromatic green or UV flash stimulation, respectively. Seven different green light stimulus intensities between 0.64 and 214 mW/sr/m^2^, and eight different UV intensities between 1.53 and 509 mW/sr/m^2^, were used in ascending order. The whole stimulation protocol lasted less than 10 min. Stimulus intensities in radiance units were obtained from measurements and conversion coefficients provided by Diagnosys LLC (Lowell, MA). The details of conversion of luminous energy units to radiance units are presented in Supplementary Table 1.

The ERG signal was acquired at 2 kHz and filtered with a low-frequency cutoff at 0.25 Hz and a high-frequency cutoff at 300 Hz. Espion software automatically detected the ERG a-wave (first negative ERG component) and b-wave (first positive ERG component) amplitudes; a-wave amplitude was measured from the signal baseline, whereas b-wave amplitude was measured as the difference between the negative trough (a-wave) and the highest positive peak. For assessment of retinal light sensitivity (I_1/2_), b-wave amplitudes were fitted as a function of stimulus intensity by using the Naka-Rushton equation (Naka and Rushton, 1966), wherein the I_1/2_ parameter describes the light intensity needed to exert half-maximal b-wave.

### Optomotor response behavioral vision test

The optomotor reflexes (OMR) were assessed using a commercial OMR platform (Phenosys qOMR, PhenoSys GmbH, Berlin, Germany) that utilizes automated head tracking and behavior analysis, following protocols described elsewhere (Suh *et al*., 2020). The OMR arena was lit at ∼ 80 lux, corresponding to the photopic light level. Rotating (12° sec^−1^) vertical sinusoidal grating stimuli at various spatial frequencies (SF) were presented for 11 min per trial to light-adapted mice. The contrast between the white and black gratings was set at 100%, whereas the SF (0.05, 0.1, 0.15, 0.20, 0.25, 0.30, 0.35, 0.375, 0.40, 0.425, 0.45 cycles per degree of visual angle, CPD) pattern changed every 60 sec in a random order, with one exception; each session always started with 0.1 CPD to facilitate acclimatization to the task, as this SF is known to evoke reliable OMR in WT mice. Each mouse was tested in at least four trials. The performances across the trials were averaged for analysis, excluding those 60-sec stimulus periods that led to a correct/incorrect ratio smaller than 0.8. All experiments were performed in the morning before noon.

### Single unit and local field potential recordings and visual stimulation

Mice were initially anesthetized with 2% isoflurane in a mixture of N_2_O/O_2_ (70%/30%), then placed into a stereotaxic apparatus. A small, custom-made plastic chamber was glued (Vetbond, St. Paul, MN, US) to the exposed skull. After one day of recovery, re-anesthetized animals were placed in a custom-made hammock, maintained under isoflurane anesthesia (1-2% in N_2_O/O_2_), and multiple single tungsten electrodes were inserted into a small craniotomy above the visual cortex. Once the electrodes were inserted, the chamber was filled with sterile agar and sealed with sterile bone wax. During recording sessions, animals were sedated with chlorprothixene hydrochloride (1 mg/kg, IM; (Camillo *et al*., 2018)) and kept under light isoflurane anesthesia (0.2 – 0.4% in 30% O_2_). EEG and EKG were monitored throughout the experiments, and body temperature was maintained with a heating pad (Harvard Apparatus, Holliston, MA).

Data were acquired using a 32-channel Scout recording system (Ripple, UT, USA). The local field potential (LFP) from multiple locations was bandpass filtered from 0.1 Hz to 250 Hz and stored together with spiking data on a computer with a 1 kHz sampling rate. The LFP signal was cut according to stimulus time stamps and averaged across trials for each recording location to calculate visually evoked potentials (VEP) (Foik *et al*., 2015; Kordecka *et al*., 2020; Suh et al., 2020; Lewandowski *et al*., 2022). The spike signal was bandpass filtered from 500 Hz to 7 kHz and stored in a computer hard drive at a 30 kHz sampling frequency. Spikes were sorted online in Trellis (Ripple, UT, USA) while performing visual stimulation. Visual stimuli were generated in Matlab (Mathworks, USA) using Psychophysics Toolbox (Brainard, 1997; Pelli, 1997; Kleiner *et al*., 2007) and displayed on a gamma-corrected LCD monitor (55 inches, 60 Hz; 1920 × 1080 pixels; 52 cd/m^2^ mean luminance). Stimulus onset times were corrected for LCD monitor delay using a photodiode and microcontroller (in-house design) (Foik *et al*., 2018).

The vision quality was assessed using protocols published in our previous work (Foik *et al*., 2018, 2020; Kordecka *et al*., 2020; Suh *et al*., 2020). For recordings of visually evoked responses, cells were first tested with 100 repetitions of a 500 msec bright flash of light (105 cd/m^2^). Receptive fields for visually responsive cells were then located using square-wave drifting gratings, after which optimal orientation/direction, spatial and temporal frequencies were determined using sine-wave gratings. SFs tested were from 0.001 to 0.5 cycles/º. Temporal frequencies tested were from 0.1 to 10 cycles/sec. With these optimal parameters, size tuning was assessed using apertures of 1 to 110º at 100% contrast. With the optimal size, temporal and SF, and at high contrast, the orientation tuning of the cell was tested again using 8 orientations x 2 directions each, stepped by 22.5º increments. This was followed by testing contrast. Single units were recorded along the whole depth of the primary visual cortex as presented in previous studies (Foik *et al*., 2018; Suh et al., 2020; Frankowski *et al*., 2021; Choi *et al*., 2022; Lewandowski *et al*., 2022).

### VEP data analysis

The response amplitude of LFP was calculated as a difference between the peak of the positive and negative components in the VEP wave. The response latency was defined as the time point where maximum response occurred. The maximum of the response was defined as the maximum of either the negative or positive peak.

Tuning curves were calculated based on the average spike rate. Optimal visual parameters were chosen as the maximum response value. Orientation tuning was measured in degrees as the half-width at half-height (HWHH; 1.18 x σ) based on fits to Gaussian distributions (Carandini and Ferster, 2000; Alitto and Usrey, 2004; Foik *et al*., 2018, 2020) using:

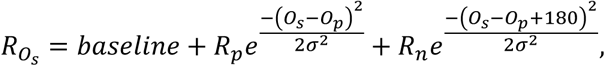

where O_s_ is the stimulus orientation, R_Os_ is the response to different orientations, O_p_ is the preferred orientation, R_p_ and R_n_ are the responses at the preferred and non-preferred direction, σ is the tuning width, and ’ baseline’ is the offset of the Gaussian distribution. Gaussian fits were estimated without subtracting spontaneous activity, similar to the procedures of Alitto and Usrey (Alitto and Usrey, 2004).

Size tuning curves were fitted by a difference of Gaussian (DoG) function:

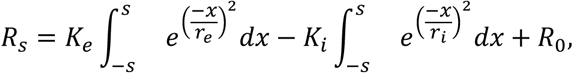

in which R_s_ is the response evoked by different aperture sizes. The free parameters, K_e_ and r_e_, describe the strength and the size of the excitatory space, respectively; K_i_ and r_i_ represent the strength and the size of the inhibitory space, respectively; and R_0_ is the spontaneous activity of the cell.

The optimal spatial and temporal frequencies were extracted from the data fitted to Gaussian distributions using the following equation (DeAngelis *et al*., 1993; Van Den Bergh *et al*., 2010; Foik *et al*., 2018, 2020):

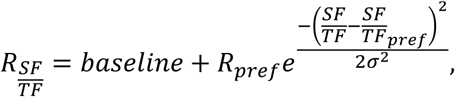

where R_SF/TF_ is the estimated response, R_pref_ indicates response at a preferred spatial or temporal frequency. SF/TF indicates spatial or temporal frequency, σ is the standard deviation of the Gaussian, and the baseline is the Gaussian offset.

The contrast tuning was fitted by using the Naka-Rushton equation (Naka and Rushton, 1966; Albrecht and Hamilton, 1982; Van Den Bergh *et al*., 2010; Przybyszewski *et al*., 2014; Foik *et al*., 2018):

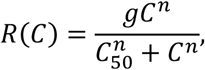

where g is the gain (response), C_50_ is the contrast at mid response, and n is the exponent. For the contrast tuning fit the background activity was subtracted from the response curve, and values below background standard deviation were changed to 0, as done elsewhere (Van Den Bergh *et al*., 2010; Foik *et al*., 2018).

### Spectral dissection of ERG and VEP responses

The ERG and VEPs were filtered using 4^th^ order Butterworth bandpass filters of 13 - 29 Hz for Beta, and 30 - 120 Hz for Gamma ranges. Two filtered signals for each animal were then used to calculate amplitude spectra using the fast fourier transform (FFT) algorithm (Matlab, USA) and amplitude spectra area (AMSA), and shown as bar plots. The dominant frequencies in beta and gamma ranges were picked for each animal separately as the frequency at maximum amplitude in the calculated spectrum. This process was repeated for each animal in each group and for green and UV light intensities to create the frequency *vs*. light intensity and AMSA *vs*. light intensity traces.

### Statistical analysis

Data throughout the manuscript are presented as mean ± SEM. The level of statistical significance was set at *P* < 0.05. The ERG light intensity series were analyzed by two-way repeated measures ANOVA (RM ANOVA), using the light intensity as the within-subjects factor (repeated) and genotype as the between-subjects factor. Sidak’s *post hoc* tests were applied if ANOVA found a significant between-subjects main effect, or within between-subjects interaction. Retinal light sensitivity (I_1/2_) was analyzed by Welch’s t-test. In the OMR data, the genotypes were compared using regular two-way ANOVA, whereas repeated measurements in the same animals (*e.g*., WT animals at 1 *vs*. 3-month of age) were analyzed using two-way RM ANOVA by age as the repeated factor. GraphPad Prism 9 (San Diego, CA) software was used for ERG and OMR data analysis. Data from cortical recordings and oscillatory activity was analyzed with the Kruskal-Wallis test followed by Dunn-Sidak *post-hoc* tests. Cortical recording and oscillatory activity offline data analysis and statistics were performed in Matlab (Mathworks, USA).

## Results

### Increased photopic ERG b-wave sensitivity in Rho^P23H/WT^ mice

We began our investigations by recording photopic ERGs in light-adapted 1-month-old mice (Figure 1A-H). Because mouse S- and M-cones are most sensitive for UV and green light, with peak sensitivities at around 360 nm and 510 nm (Jacobs *et al*., 1991; Lyubarsky *et al*., 1999), respectively, and since S- and M-cones could be differentially affected during retinal degeneration (Greenstein et al., 1989; Schwarz *et al*., 2018; Hassall *et al*., 2020), we recorded photopic ERGs using both monochromatic UV light and green light stimuli. The a-or b-wave response amplitudes did not show an overall difference between genotypes as determined by repeated measures analysis of variance (RM ANOVA; a-wave, Supplementary Figure 1A,B; b-wave, Figure 1B,F). However, anecdotally *Rho*^P23H/WT^ mice seemed to display higher amplitudes at intermediate light intensities (Figure 1B,F). Therefore, we used the Naka-Rushton function for curve fitting (Naka and Rushton, 1966) and derivatization of light intensity required to generate half-maximal b-wave responses (I_1/2_) as a measure of retinal light sensitivity. We found decreased I_1/2_, indicative of increased sensitivity, in *Rho*^P23H/WT^ mice for both green light (Figure 1C) and UV light (Figure 1G) stimulation, which supports previous findings (Leinonen *et al*., 2020). In addition, b-wave peak latency on average was faster in *Rho*^P23H/WT^ mice as compared to WT mice (Figure 1A, H), but this trend towards shorter latency did not reach a statistically significant value (green, *P*=0.09; UV, *P*=0.20). Instead, the a-wave latency was significantly faster in response to a UV flash in 1-month-old *Rho*^P23H/WT^ mice compared to WT (Supplementary Figure 1F), but not in response to a green flash (Supplementary Figure 1E, *P*=0.73).

**Figure 1.**
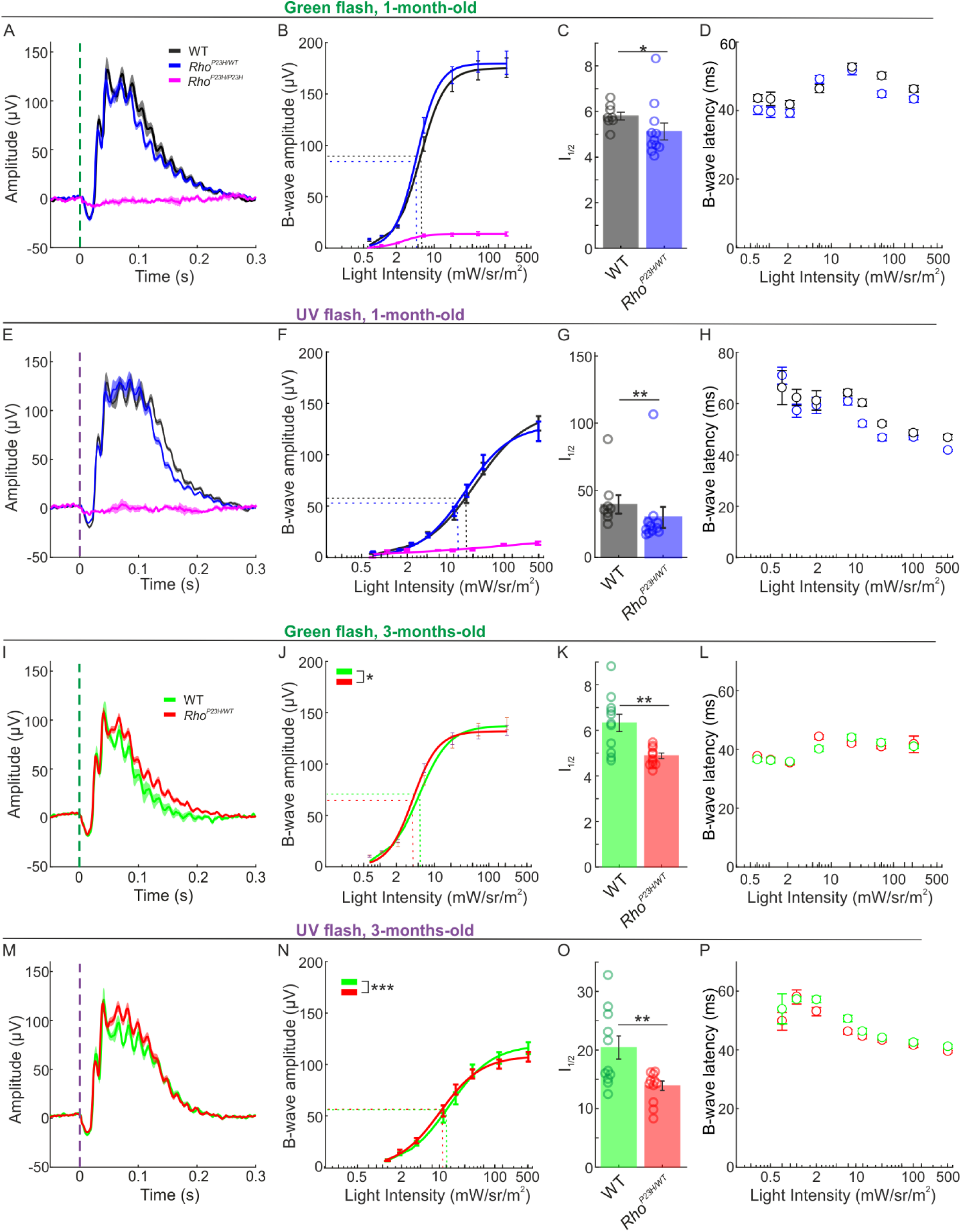
Increased light sensitivity in cone-dominant ERGs in Rho^P23H/WT^ mice. Panels A-H present data from 1-month-old mice, and panels I-P correspond to 3-month-old mice. (A) ERG waveforms in response to a 214 mW·sr/m^2^ green light flash (peak emission 544 nm, bandwidth 160 nm; black/grey lines, WT, n=8; blue, *Rho*^*P23H/WT*^, n=11; magenta, *Rho*^*P23H/P23H*^, n=4). Lines represent group averages ± SEM. (B) b-wave amplitude series and (C) b-wave sensitivity (I_1/2_) to green light flash. (D) b-wave latency series to green flash. (E) ERG waveforms in response to a 509 mW·sr/m^2^ UV flash (peak emission 370 nm, bandwidth 50 nm). (F) b-wave amplitude series. (G) b-wave sensitivity (I_1/2_) to UV light flash. (H) b-wave latency series. (I) ERG waveforms in response to a 214 mW·sr/m^2^ green flash in 3-month-old mice (green lines, WT, n=11; red, *Rho*^*P23H/WT*^, n=11). (J) b-wave amplitude series (light intensity-response amplitude interaction, *P*<0.05). (K) I_1/2_ for green light flash. (L) b-wave latency series for green flash. (M) ERG waveforms in response to 509 mW·s/m^2^ UV flash in 3-month-old mice. (N) b-wave amplitude series for UV flashes (light intensity-response amplitude interaction, *P*<0.001). (O) I_1/2_ for UV light flash. (P) b-wave latency series for UV flash. Repeated measures two-way ANOVA were applied to amplitude and latency analyses in the increasing flash intensity series. I_1/2_ was acquired from curve fitting, following the Naka-Rushton equation and derivation of light intensity required to generate a half-maximal response. Comparisons for the I_1/2_ parameter were performed by Welch’ s t-test. Data are presented as mean ± SEM, plus individual replicates (circles) in C, G, K, and O.

To study whether increased photopic ERG sensitivity is persistent in adulthood, we performed the same recordings at 3-months of age (Figure 1I-P). Here, the maximal b-wave amplitudes in *Rho*^P23H/WT^ mice tended to remain slightly lower compared to WT (Figure 1 J, N), although no significant RM ANOVA effects were found for the amplitude comparisons (green, *P*=0.25; UV, *P*=0.76). Instead, RM ANOVA revealed a significant light intensity-response amplitude interaction both in response to green light (Figure 1J, *P*<0.05) and UV light (Figure 1N, *P*<0.001) stimulation. The RM ANOVA interaction together with visual inspection of light intensity-response amplitude curves (Figure 1J,N) indicate that although maximal b-wave amplitudes tended to decrease in *Rho*^P23H/WT^ mice, responses to intermediate light intensities were above WT level. Indeed, *Rho*^P23H/WT^ mice still showed significantly decreased I_1/2_ b-wave sensitivity parameters compared to WT mice (Figure 1K, O), regardless of intermediate retinal degeneration at this stage (Sakami *et al*., 2011; Leinonen *et al*., 2020). We interpret the relative decrease in b-amplitude and increase in b-wave latency in *Rho*^P23H/WT^ mice between 1- and 3-months of age (compare Figures 1B, D, F, H to Figures 1J, L, N, P) to reflect a disease stage when cone degeneration is commencing, although there were not yet decreases in a-wave amplitudes at this age range (Supplementary Figure 1C, D). Overall, the modest photopic ERG changes which we observed in *Rho*^P23H/WT^ mice appeared to be more evident in the S-cone pathway as compared to the M-cone pathway.

To provide a context for how the advanced-disease stage represents a cone-mediated retinal function, we recorded photopic ERGs also in 1-month-old homozygous mutant (*Rho*^P23H/P23H^) mice with an almost complete loss of photoreceptors by this age (Sakami *et al*., 2011). These recordings demonstrated only a residual response in the cone-pathway (Figure 1A, B, E, F & Supplementary Figure 2), which rendered curve fitting, light sensitivity, and response latency assessments impractical.

Prominent oscillatory potentials (OPs) ride on the ascending phase of the rodent ERG b-wave and could have implications on the b-wave analyses. We therefore performed OP amplitude and peak frequency analyses at two distinct ranges: beta (13 – 29 Hz) and gamma (30 – 120 Hz) (Supplementary Figure 4). The OP amplitude was significantly attenuated in Rho^P23H/P23H^ mice as expected, due to extreme retinal degeneration and resulting minimal input from photoreceptors. Instead, there were no significant differences between the *Rho*^P23H/WT^ and WT mice either in AMSA or dominant frequency parameter at any of the conditions (Supplementary Figure 4), except for one: *Rho*^P23H/WT^ mice displayed a higher dominant frequency than WT mice in response to the strongest UV flash at the beta range (Supplementary Figure 4J,K). Overall, OPs cannot explain the increased b-wave sensitivity observed in *Rho*^P23H/WT^ mice (Figure 1).

### Rho^P23H/WT^ mice display increased optomotor movements towards low spatial frequency (SF) drifting patterns

We next sought to determine if visual acuity, an elementary visual behavior as measured by optomotor responses, is changed in light-adapted/photopic conditions in 1.5 and 3-month-old *Rho*^P23H/WT^ mice. An earlier report using OMR found contrast sensitivity was well-preserved in *Rho*^P23H/WT^ mice even in scotopic conditions despite prominent rod death (Leinonen *et al*., 2020). Here we utilized a commercial OMR setup coupled with an automated head-tracking system (Kretschmer *et al*., 2013, 2015), which allowed us to record the mouse optomotor movements with respect to the drifting pattern in its visual field at 30 frames per second. The common OMR studies have typically investigated the SF (visual acuity assessment) or contrast threshold detection limits using subjective assessment. In contrast, our paradigm ranks correct/incorrect optomotor movements at every frame, providing a more sensitive and objective means to document ” supernormal” visual behavior. At the highest SF (smallest pattern sizes) the stimuli elicited similar responses in the 1.5-month-old *Rho*^P23H/WT^ mice and WT mice; however, with the *Rho*^P23H/WT^ mice, optomotor movements towards stimuli at the lowest SF (largest pattern size) were more frequent than those of WT mice, as demonstrated by their higher OMR index (Figure 2A). This behavior reflects sensorimotor hypersensitivity to low SF drifting patterns in the visual field of the *Rho*^P23H/WT^ mice. The same mice performed the same task later at 3 months of age. Head-tracking behavior in *Rho*^P23H/WT^ mice slightly decreased between 1.5 and 3 months of age (Figure 2B), indicating an initial decline in visual acuity in 3-month-old *Rho*^P23H/WT^ mice. Still, *Rho*^P23H/WT^ mice at 3 months of age continued with a supernormal OMR at low SF, as compared to same-age WT mice (Figure 2C; OMR index-SF interaction, *P*<0.05). Intact responses in WT mice between 1.5 and 3 months of age document the feasibility of the repeated analysis (Figure 2D, *P* = 0.49).

**Figure 2.**
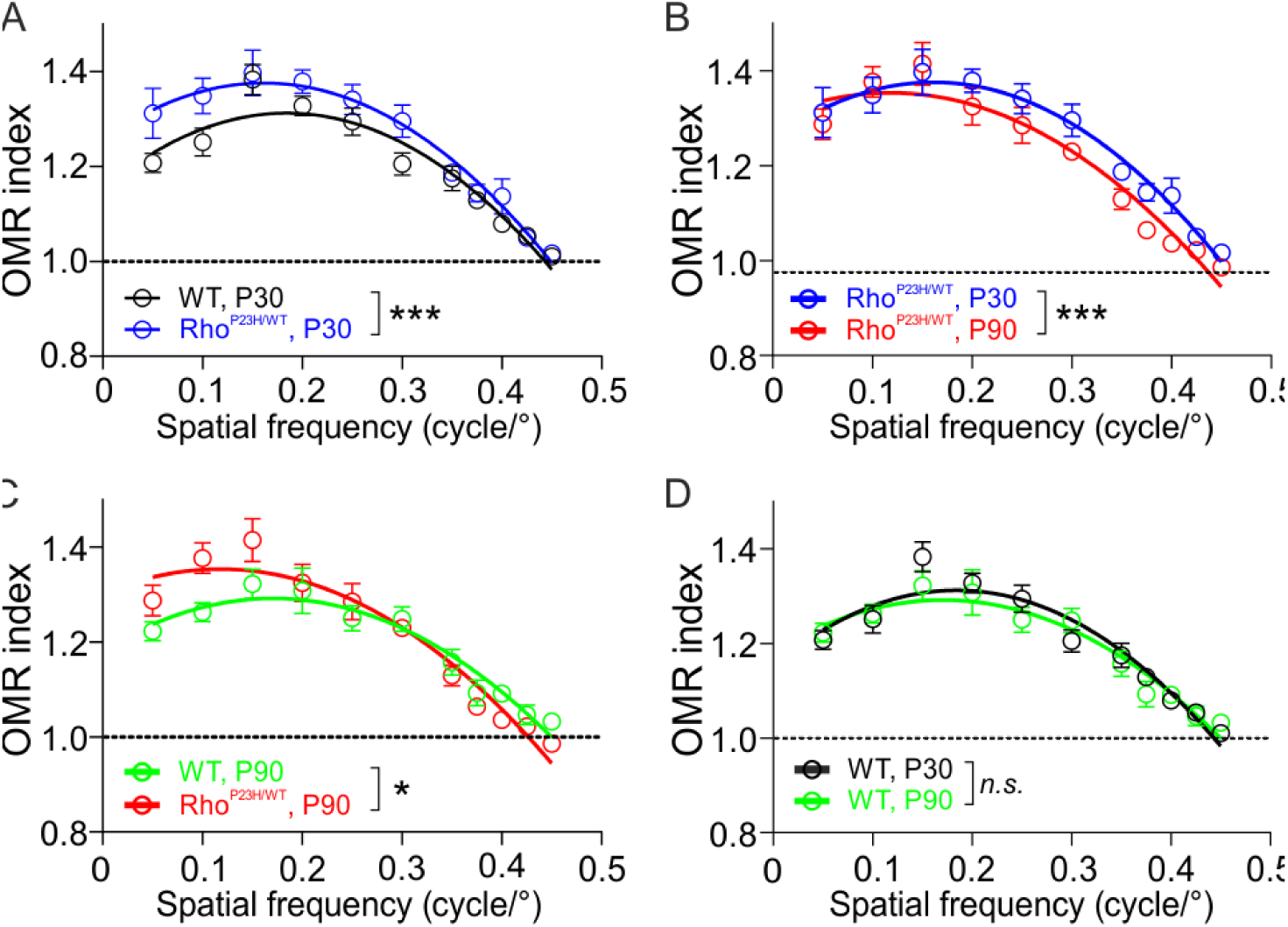
Hypersensitized optomotor response (OMR) to low spatial frequency (SF) drifting gratings in Rho^P23H/WT^ mice. (A) Group comparison of WT (n=8) and *Rho*^P23H/WT^ (n=10) mice at 1.5-months of age (genotype effect: \*\*\**P*<0.001). (B) Repeated analysis in *Rho*^*P23H/WT*^ mice at 1.5 and 3 months of age (age effect: \*\*\**P*<0.001). (C) Group comparison of WT (n=8) and *Rho*^P23H/WT^ (n=10) mice at 3-months of age (OMR index-SF interaction: \**P*<0.05). (D) Repeated analysis in WT mice at 1.5 and 3 months of age (age effect: *P*=0.49). The statistical analysis was performed using two-way RM ANOVA. OMR index-SF interaction in C signifies opposing sensitivity differences between WT and *Rho*^P23H/WT^ groups at low versus high SF. n.s., nonsignificant. Data are presented as mean ± SEM. Lines that represent second-order quadratic polynomial fits are for illustration and were not used in the analysis.

### Stimulus-dependent and spontaneous visual cortex hyperexcitability in juvenile Rho^P23H/WT^ mice

Most importantly for central vision, it was important to investigate the consequences of early retinal degeneration for light responses at the V1 level (note, OMR is a subcortically originating reflex and does not require primary visual cortex (V1) function (Douglas *et al*., 2005)). To this end, we started by recording visual evoked potentials (VEPs) in response to light ON/light OFF stimuli. We found an increase in both ON and OFF responses in 1-month-old *Rho*^P23H/WT^ mice compared to same-age WT mice (Figure 3A, B, C, E), indicating hyperexcitability. Conversely, the same age homozygous *Rho*^P23H/P23H^ mutants showed only a residual response as expected, due to severe retinal degeneration and dysfunction (Figure 3C,E; (Sakami *et al*., 2011)). Heterozygous *Rho*^P23H/WT^ mice at 3 months old, when more than half of their photoreceptors are gone (Leinonen *et al*., 2020), still showed VEP amplitude comparable to WT mice (Figure 3C, E). The response latency to the ON stimulus increased with disease severity (Figure 3D, *P<*0.01), whereas latency to the OFF stimulus remained intact in heterozygous mutants (Figure 3F).

**Figure 3.**
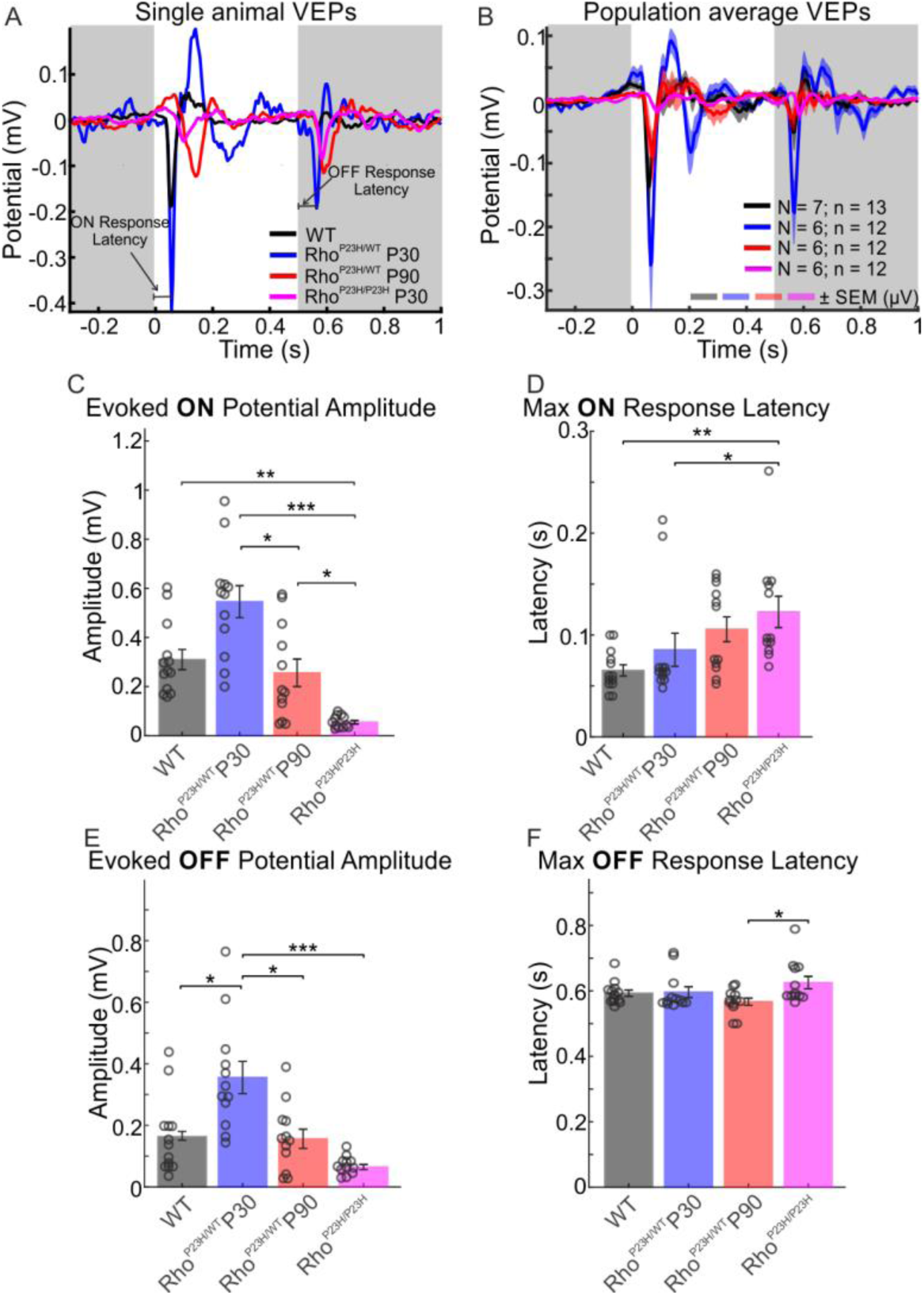
Supernormal visual evoked potentials (VEPs) for light ON/light OFF stimuli in juvenile Rho^P23H/WT^ mice. White and gray backgrounds in A and B indicate the period when the light was ON or OFF, respectively. (A) Representative VEP waveforms from a single WT mouse (black), a 1-month-old *Rho*^P23H/WT^ mouse (blue line), a 3-month-old *Rho*^P23H/WT^ mouse (red), and a 1-month-old *Rho*^P23H/P23H^ mouse (magenta). Horizontal lines indicate ON-response latency or OFF-response latency, as indicated. (B) Group-averaged VEP waveforms (WT, N = 7 mice, n = 13 eyes; 1-month-old *Rho*^P23H/WT^, N = 6 mice, n = 12 eyes; 3-month-old *Rho*^P23H/WT^, N = 6 mice, n = 12 eyes; 1-month-old *Rho*^P23H/P23H^, N = 6 mice, n = 12 eyes). The colored shading in B indicates ± SEM from the response mean. (C) ON response amplitudes. (D) ON response latencies. (E) OFF response amplitudes. (F) OFF response latencies. Data are presented as mean ± SEM, plus individual replicates (circles) in C-F. The statistical analysis was performed using the Kruskal-Wallis test followed by Dunn-Sidak post hoc tests: \**P*<0.05, \*\**P*<0.01, \*\*\**P*<0.001.

To investigate if oscillatory activity is also affected in V1 in response to ON and OFF stimuli, we analyzed AMSA as well as dominant oscillation frequency at beta (13-29 Hz), and gamma frequency (30-120 Hz) ranges (Figure 4A, B and Figure 4E, F, respectively). As with the main VEP component amplitudes, we found on average higher AMSAs in the beta (Figure 4B, C) and gamma frequency (Figure 4F, G) ranges in 1-month-old *Rho*^*P23H/WT*^ mice, although far from statistical significance (*P*=0.33 and *P*=0.46, respectively). Instead, by 3-months of age, the amplitude of oscillatory activity lowered substantially in *Rho*^*P23H/WT*^ mice and reached levels similar to WT (Figure 4B, C; P=0.79). As expected, AMSAs at both beta and gamma ranges were significantly lower in the most advanced disease stage studied, in *Rho*^P23H/P23H^ mice (Figure 4B, C, F, G). We found no differences in dominant oscillation ranges at either the beta (Figure 4D, *P*=0.13) or gamma ranges (Figure 4H, *P*=0.3). Overall, the lack of major differences in dominant oscillatory frequencies suggests no change in large network processing in the brain.

**Figure 4.**
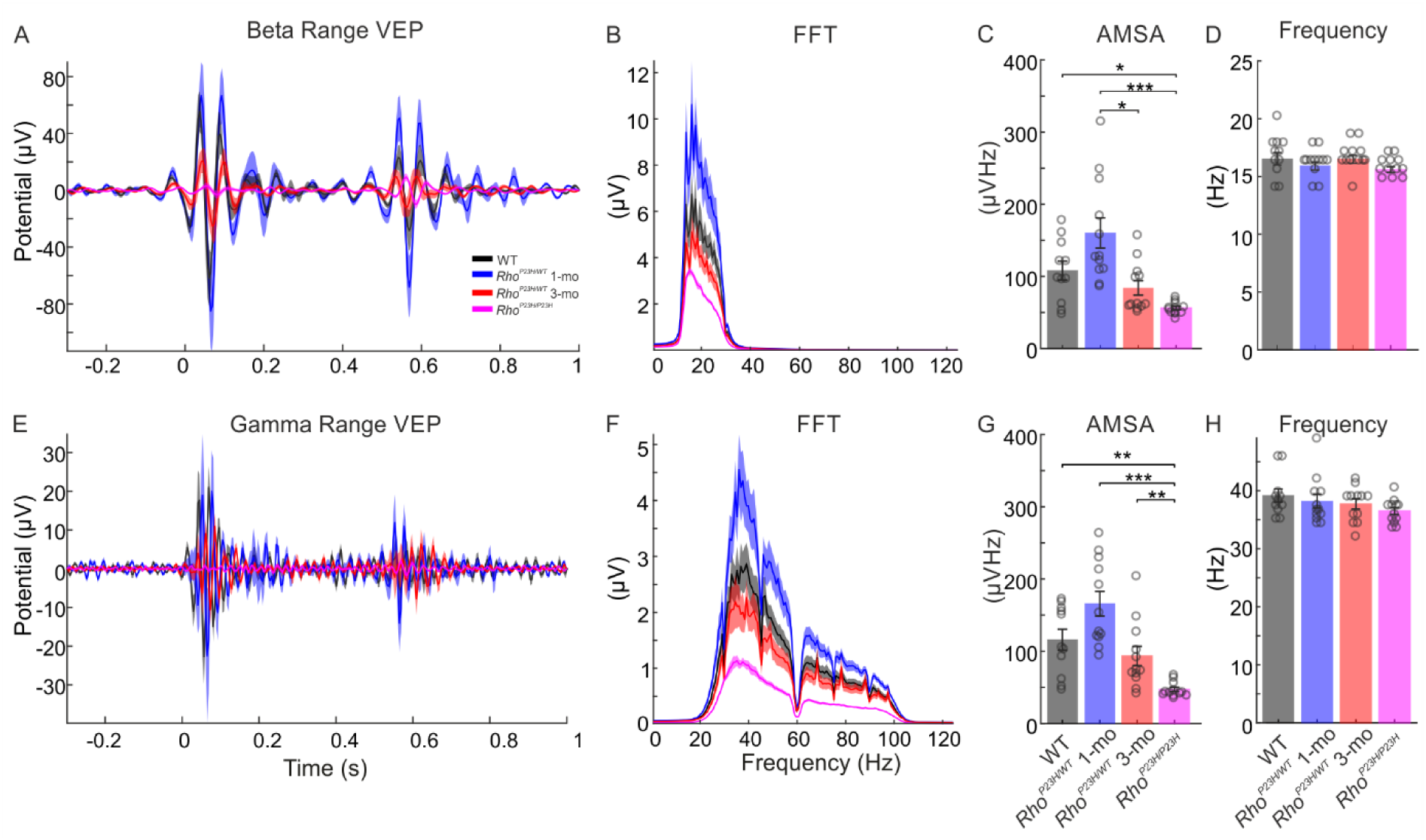
Increased oscillatory activity in response to light in V1 of juvenile Rho^P23H/WT^ mice. Beta and gamma range oscillations in VEP responses to light flash are shown. (A) Average cortical VEPs for all animal groups were filtered using the bandpass filter in the beta frequency range (13 - 29 Hz). (B) Corresponding fast fourier transform (FFT) spectra are shown. (C) Comparison of population-average amplitude spectrum areas (AMSA) in the beta frequency range. (D) Comparison of population-average dominant oscillation frequencies in the beta frequency range. (E) Average cortical VEPs for all groups were filtered using the bandpass filter in the gamma range (30 - 120 Hz). (F) Amplitude spectra were calculated for each animal group. Comparison of AMSA values among all groups in the gamma frequency range. (H) Population-average dominant frequencies of oscillations in the gamma range. Circles indicate individual recording sites. The Kruskal-Wallis test was performed, followed by Dunn-Sidak post hoc tests: *P<0.05, **P<0.01, ***P<0.001.

Single-unit recordings corroborated significant visual cortex hyperexcitability in the juvenile *Rho*^P23H/WT^ mice (Figure 5). Both light ON and light OFF conversions evoked more spiking activity in 1-month-old *Rho*^P23H/WT^ mice relative to WT (Figure 5A, B, G, H). One should note, however, that more than half of the V1 neurons in *Rho*^P23H/WT^ mice did not respond to the light stimulus, whereas in WT mice less than 10% of units were non-responsive (Figure 5E). The cortical hyperexcitability in juvenile *Rho*^P23H/WT^ mice was not only stimulus-dependent but also continuous as the stimulus-independent background spiking was also significantly elevated in the juvenile *Rho*^P23H/WT^ mice *versus* WT (Figure 5A, B, F). The spiking activity diminished in the mutant mice with disease progression, so that average spiking activity to light ON or OFF did not differ between 3-month-old *Rho*^P23H/WT^ and WT mice (Figure 5A, C, G, H). Notably, however, light ON responses in *Rho*^P23H/WT^ mutants were qualitatively abnormal regardless of age, as highlighted by the second peak in spiking activity occurring at ∼ 200 ms or later from light onset (Figure 5C), a phenomenon that was absent in the WT mice (Figure 5A). The spiking pattern at light OFF remained relatively normal in *Rho*^P23H/WT^ mutants (Figure 5A-C). As expected, the spiking activity to both light ON or light OFF was significantly dampened in the homozygous *Rho*^P23H/P23H^ mice (Figure 5D, G, H). Conversely, the spontaneous spiking activity was *increased* in late-stage disease (*Rho*^P23H/P23H^) compared to the less advanced disease stage (3-month-old *Rho*^P23H/WT^) (Figure 5F), indicating that some phenomenon independent of photoreceptors begins to drive spontaneous V1 neuron spiking in late-stage retinal degeneration, similar to what has been proposed previously regarding activity recordings of cone bipolar, amacrine, and retinal ganglion cells (Borowska *et al*., 2011; Menzler and Zeck, 2011).

**Figure 5.**
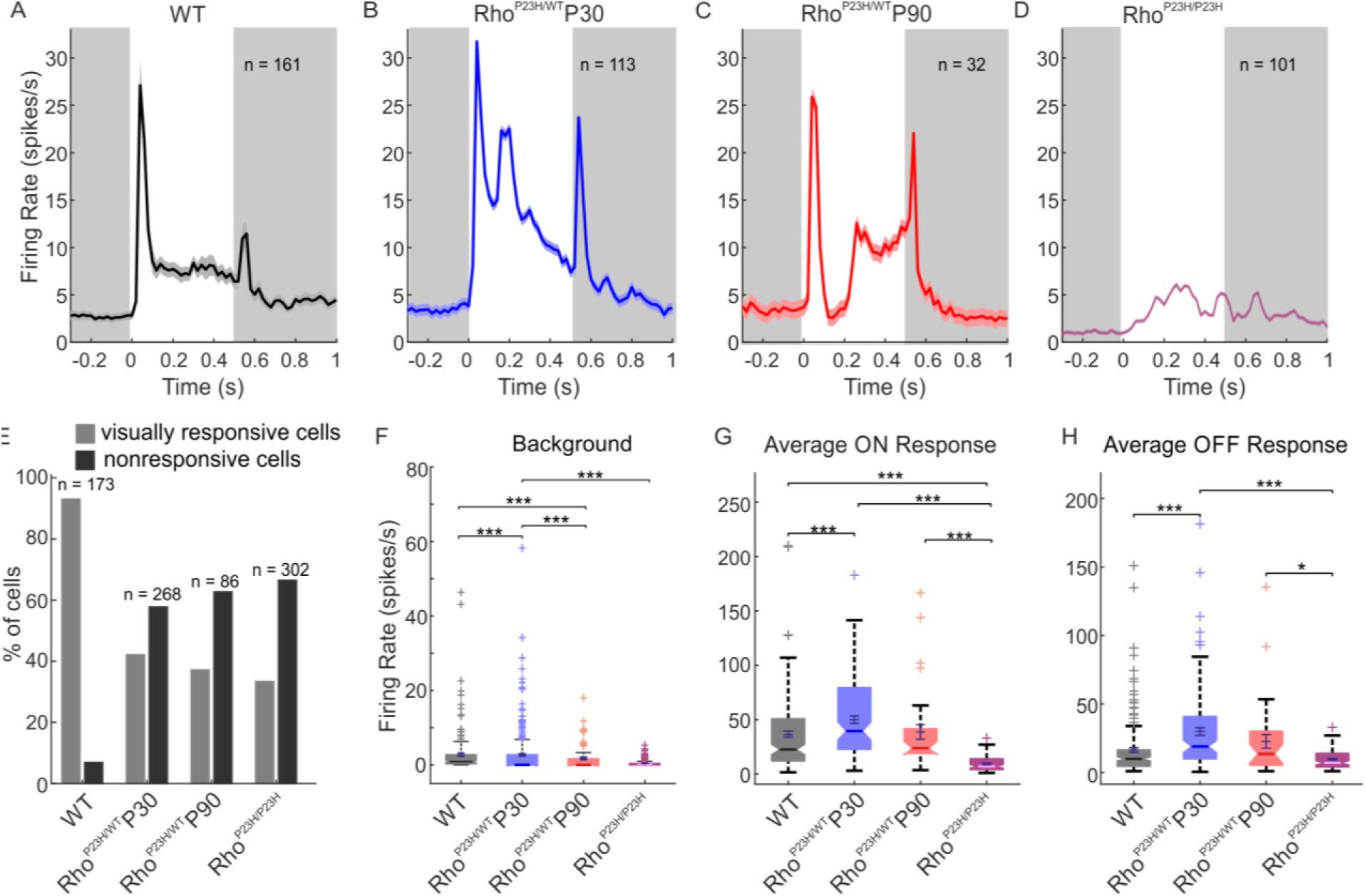
Cortical neurons are hyperexcited in juvenile Rho^P23H/WT^ mice. White and gray backgrounds in A and B indicate the period when the light was ON or OFF, respectively. (A) Population summary of single-neuron responses to light ON/OFF stimuli for WT mice (N = 7 mice, n = 161 neurons); (B) 1-month-old *Rho*^P23H/WT^ mice (N = 6 mice, n = 113 neurons); (C) 3-month-old *Rho*^P23H/WT^ mice (N = 6 mice, n = 32 neurons); and (D) 1-month-old *Rho*^P23H/P23H^ mice (N = 6 mice, n = 101 neurons). (E) Percentage of visually responsive and non-responsive neurons in each group. (F) Background firing rate without light stimulus. (G) Maximum firing rate at light ON for each group. (H) Maximum firing rate at light OFF for each group. Each box plot (F-H) indicates a median marked by a solid line and a notch. The bottom and top edges of the boxes indicate 25^th^ and 75^th^ percentiles, and whiskers indicate maximum and minimum values in distributions. Crosses indicate individual values treated as outliers. Kruskal-Wallis test followed by Dunn-Sidak post hoc tests: \**P*<0.05, \*\**P*<0.01, \*\*\**P*<0.001.

### Compromised tuning properties of V1 neurons to drifting pattern stimuli

We last investigated V1 neuron tuning properties in *Rho*^P23H/WT^ mice. For this task, we used drifting sinusoidal-pattern stimuli to study several different parameters relevant for visual behavior: orientation/direction tuning, SF tuning, as well as thresholds for SF, contrast, and temporal frequency. Several abnormalities were observed in *Rho*^P23H/WT^ mice. First, the orientation tuning curve was distinctly broader compared to WT mice already at 1-month of age (Figure 6A, B). Strikingly, the 3-month-old *Rho*^P23H/WT^ mice practically preferred no directions, demonstrating a dramatic loss of directional selectivity. Next, the pattern size to which mice optimally responded increased with disease progression, indicating enlargement of receptive fields (Figure 6C, D). The optimal SF was slightly decreased at 1-month-old for *Rho*^P23H/WT^ mice, and severely decreased by the age of 3-months (Figure 6E, F). In contrast, the optimal temporal frequency was abnormally high in 1-month-old *Rho*^P23H/WT^ mice (Figure 6G, H), mirroring hastened kinetics of retinal responses to light stimulation (Figure 1D, H, and Supplementary Figure 1F). Also, the contrast sensitivity in *Rho*^P23H/WT^ mice remained at similar levels to those for WT mice, even at 3 months of age (Figure 6I, J). Altogether, the VEP and V1 single-unit recordings observed in young *Rho*^P23H/WT^ mice reveal that hastened ERG kinetics (Figure 1D, H; Supplementary Figure 1F; (Pasquale *et al*., 2021)), maintained behavioral contrast and temporal frequency thresholds (Leinonen *et al*., 2020; Pasquale *et al*., 2021), and increased preference for lower SF stimuli (Figure 2A, C) are reflected in V1 function. We were unable to record meaningful responses to pattern stimuli in the most advanced disease state, in *Rho*^P23H/P23H^ mice.

**Figure 6.**
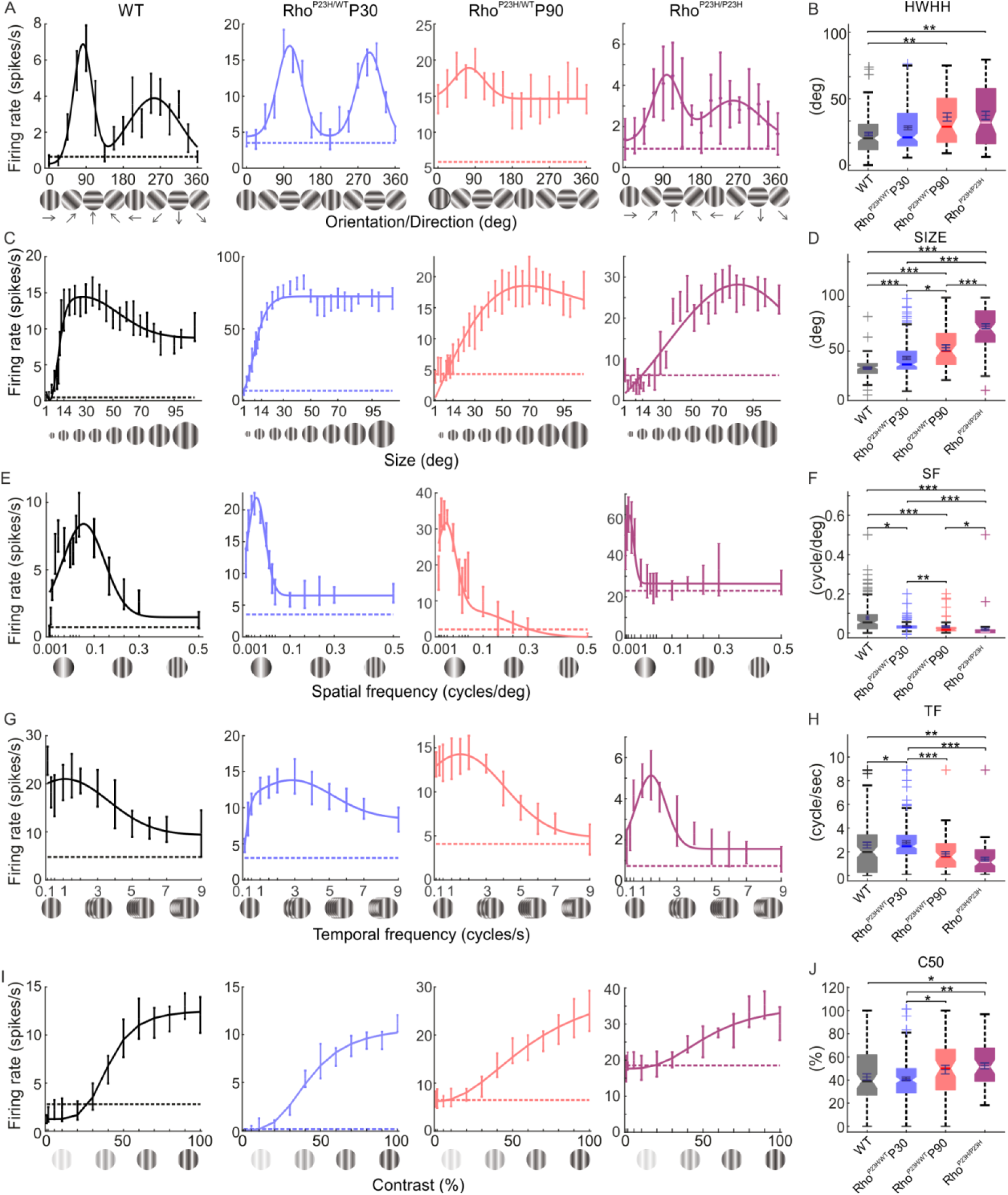
Altered V1 neuron-tuning properties in Rho^P23H/WT^ mice. (A) Representative orientation/direction tuning curves of single cells for all groups. (B) Orientation tuning is measured as half-width at half-height (HWHH) (WT, black, N = 7 mice, n = 153 neurons; 1-month-old *Rho*^P23H/WT^, blue, N = 6 mice, n = 189 neurons; 3-month-old *Rho*^P23H/WT^, blue, N = 6 mice, n = 116 neurons; 1-month-old *Rho*^P23H/P23H^, magenta, N = 6 mice, n = 88 neurons). (C) Representative stimulus size tuning curves of single cells. (D) Stimulus size at which maximal firing rate was observed. (E) Representative spatial frequency (SF) tuning curves of single cells. (F) SF at which maximal firing rate was observed. (G) Representative temporal frequency tuning curves of single cells. (H) Drifting pattern speed at which maximal firing rate was observed. (I) Representative contrast tuning curves of single cells. (J) Pattern contrast yielded half of the maximum response (C50). Each box plot indicates a median marked by a solid line and a notch. The bottom and top edges of the boxes indicate 25^th^ and 75^th^ percentiles, and whiskers indicate maximum and minimum values in distributions. Crosses indicate individual values treated as outliers. Statistical analysis was performed using the Kruskal-Wallis test followed by Dunn-Sidak post hoc tests: \**P*<0.05, \*\**P*<0.01, \*\*\**P*<0.001.

## Discussion

### Consequences of early rod degeneration to photopic visual function

Previous studies have suggested paradoxical sensitization of cone-mediated retinal function in the earliest stages of rod degeneration in mice (Leinonen *et al*., 2017, 2020). To investigate this phenomenon further, we recorded photopic ERG responses in *Rho*^P23H/WT^ mice at 1 and 3-months of age, when ∼ 20% and ∼ 60% of rods have degenerated, respectively; but cone count remains normal (Leinonen *et al*., 2020). Maximal photopic ERG responses were intact in 1-month-old *Rho*^P23H/WT^ mice, indicating a lack of cone degeneration at this age (Figure 1). Instead, the sensitivity of the b-wave *was enhanced* compared to WT controls. At 3-months of age, the increase in b-wave sensitivity became even more apparent in *Rho*^P23H/WT^ mice despite their maximal b-wave amplitude showing some trend towards a decline (Figure 1). In a more advanced stage of RP, the cone function undoubtedly declines (Figure 1; (Sakami *et al*., 2011; Leinonen *et al*., 2020)). Increased b-wave sensitivity in rod-specific function in *Rho*^P23H/WT^/*Gnat2*^-/-^ mice have been recently observed, regardless of prominent rod degeneration (Leinonen *et al*., 2020). Relatively better-preserved ERG b-waves as compared to a-waves have been reported also in P23H transgenic rats (Machida *et al*., 2000; Aleman *et al*., 2001). The reason for increased sensitivity of the ERG b-wave is unknown but may include overcompensation of the gain at the photoreceptor-to-bipolar synapse in response to outer retina degeneration.

The kinetics of photopic ERG b-wave responses showed a trend (not statistically significant) towards hastening in 1-month-old *Rho*^P23H/WT^ mice (Figure 1). The a-wave response was significantly hastened in *Rho*^P23H/WT^ mice in response to UV stimulation (Supplementary Figure 1). Pasquale *et al*. recently reported significantly improved temporal contrast sensitivity (TCS) in *Rho*^P23H/WT^ mice (Pasquale *et al*., 2021). The improvement arose from the rod-pathway, as the phenomenon remained when *Rho*^P23H/WT^ mice were crossed with GNAT2^cpfl3/cpfl3^ mice that lack cone function. This could be explained by faster photo-response kinetics and decreased density and collecting area of rods in *Rho*^P23H/WT^ mice (Sakami *et al*., 2014; Pasquale *et al*., 2021). The mechanism that explains faster initial rod response is unclear, but could be related to volume changes in the rod outer segment or changes in Ca^2+^ dynamics (Pasquale *et al*., 2021). Our recent transcriptomic profiling of *Rho*^P23H/WT^ retina homogenates supports the latter, as we showed significant pathway enrichments in, *e.g*., calmodulin binding and ion-channel-activity-related genes, including cation channel activities (Leinonen *et al*., 2020). However, analysis at single-cell resolution is needed to test whether these changes occur in the rods or elsewhere in the retina.

As no clear deterioration of photopic ERG was observed in *Rho*^P23H/WT^ mice up to 3-months of age, and since scotopic contrast sensitivity is remarkably resistant to early rod degeneration (Leinonen *et al*., 2020), we were curious to see if any visual acuity change by OMR measures would occur at this disease stage. To this end, we used an OMR system coupled with automated head-tracking (Kretschmer *et al*., 2015). During the 60-sec recording period per SF, the system recorded 1800 frames, and at every frame compared optomotor movements to correct and incorrect directions, rendering an optomotor index. Since the OMR is a reflex-based visual task, we hypothesized that increased tracking towards the drifting patterns (or increased OMR index) by light-adapted *Rho*^P23H/WT^ mice was possible. This was partially true, as the OMR index at low SF for the *Rho*^P23H/WT^ mouse was higher compared to that for the WT mouse (Figure 2). At high SF, the OMR index was equal between genotypes at 1.5 months, indicating intact visual acuity in *Rho*^P23H/WT^ mice. However, between 1.5 and 3 months of age, the OMR index at high SF slightly declined in *Rho*^P23H/WT^ mice, indicating an initial decrease in visual acuity by 3 months in *Rho*^P23H/WT^ mice. With respect to rod-driven night vision, a strong link between enhanced scotopic ERG b-wave sensitivity and well-maintained OMR in *Rho*^P23H/WT^ and *Rho*^P23H/WT^/*Gnat2*^*-/-*^ mice was recently demonstrated (Leinonen *et al*., 2020).

### Consequences of early rod degeneration to higher-order visual processing

The increased ERG b-wave sensitivity and remarkably well-preserved pattern-contrast sensitivity (Leinonen *et al*., 2020) point to adaptive functional re-arrangements in the *Rho*^P23H/WT^ mouse retina during rod degeneration. But what happens at the primary visual cortex (V1) where the ” conscious vision” is initially processed? To address this question, we recorded VEPs and single-cell responses in V1 to light ON/light OFF and drifting pattern stimuli in lightly anesthetized mice. First, as recent OMR data have already suggested (Leinonen *et al*., 2020), pattern contrast sensitivity remained excellent in *Rho*^P23H/WT^ mice, and strikingly well-maintained even in the homozygous *Rho*^P23H/P23H^ mice with extreme retinal degeneration (Figure 6I). A corresponding persistent pattern-contrast sensitivity has been demonstrated by electrophysiology of the dorsal lateral geniculate (dLGN) nucleus in 3-5-week-old *Pde6β*^Rd1^ mice (Procyk *et al*., 2019). The *Pde6β*^Rd1^ mice at that age represent a late-stage retinal degeneration and have only a residual ERG response remaining (Strettoi *et al*., 2002). Similarly, the temporal-frequency threshold remained excellent in *Rho*^P23H/WT^ mice, and was in fact elevated at 1-month of age (Figure 6G, H). These data correspond well with the ERG (improved kinetics, Figure 1 and Supplementary Figure 1; improved TCS by ERGs; (Pasquale *et al*., 2021)) and OMR findings in *Rho*^P23H/WT^ mice (improved TCS by OMR) (Pasquale *et al*., 2021). Nevertheless, we also found detrimental changes for V1 function starting already at 1-month of age in *Rho*^P23H/WT^ mice (Figures 5 and 6). The light responses and spontaneous neuron firing were dramatically hyperexcited in the *Rho*^P23H/WT^ mouse V1 (Figures 3 and 5). The observed hyperexcitability is well in line with previous literature, demonstrating increased neural noise originating from the inner retina (Borowska *et al*., 2011; Menzler and Zeck, 2011; Trenholm and Awatramani, 2015; Haselier *et al*., 2017; Telias *et al*., 2020). Further, as similarly indicated by the OMR behavior (Figure 2), *Rho*^P23H/WT^ mouse V1 neurons preferred lower SFs compared to WT mice (Figure 6E), indicating increasing retinal ganglion cell (RGC) receptive fields. However, the most striking abnormality we found was the severely debilitated orientation/direction tuning in *Rho*^P23H/WT^ mice, which advanced to a practical absence of any direction preference by 3 months of age (Figure 6A). It is challenging to speculate how these findings translate into clinical settings, due to the lack of corresponding data from RP patients. Hypersensitivity to bright light (called by the debated misnomer ” photophobia”) is common in RP (Hamel, 2006; Otsuka *et al*., 2020), and could in part result from similar hyperexcitation of the primary visual tract as observed in RP model mice. The relatively well-maintained behavioral vision of *Rho*^P23H/WT^ mice may also be reflected in clinical settings, as many RP patients show minimal disturbances in subjective vision even at an advanced disease stage (Hartong et al., 2006).

### Relevance to treatments in retinal degenerations

There are no pharmacotherapies available that would slow retinal degenerative diseases such as RP or dry age-related macular degeneration (Waugh *et al*., 2018). It is probable, however, that the impaired photoreception in these diseases can be augmented in the near future *via* novel therapies, such as photoreceptor transplantation (Pearson *et al*., 2012), chemical photo-switches (Tochitsky *et al*., 2018), optogenetic tools (Sahel *et al*., 2021), prosthetics (Lorach *et al*., 2015), stem cells (da Cruz *et al*., 2018; Foik *et al*., 2018), or gene therapy (Suh *et al*., 2020). But a great deal of concern has been raised whether, after blinding, the more complex inner retina is capable of normal signal processing and whether restored signals are adequate for sight. Second, most patients experience only a partial loss of vision and are not suitable for advanced vision restoration therapies; therefore, more conventional approaches are needed. Using the *Pde6β*^Rd10^ mouse model of RP, Telias *et al*. recently showed that the firing rate of RGCs spontaneously increased simultaneously with decreasing light responses after mouse eye-opening in multi-electrode array recordings (Telias *et al*., 2019). They were able to link the pathology with deteriorated visual behavior in a light avoidance test by decreasing spontaneous firing pharmacologically. Importantly, blocking the retinoic acid receptor decreased spontaneous RGC firing and increased visual behavior in *Pde6β*^Rd10^ mice (Telias *et al*., 2019, 2022). Another group used a different pharmacological strategy, a gap junction blockade, but showed similar beneficial effects, wherein a decrease in spontaneous firing of RGCs led to increased light responses (Toychiev *et al*., 2013). However, the speed of degeneration is faster and the mechanism of rod death is different in the *Pde6β*^Rd10^ mouse model as compared to the *Rho*^P23H/WT^ mouse (Chang *et al*., 2007; Sakami *et al*., 2011), rendering direct comparisons difficult. It will be important in future experiments to test if decreasing the noise signal of RGCs would be therapeutic also in other models of RP, including the *Rho*^P23H/WT^ mouse with hyperexcited V1 (Figures 3-6).

On the other hand, some recent studies have concluded positive/adaptive aspects of retinal reorganization after injury or during degeneration (Beier *et al*., 2017, 2018; Johnson *et al*., 2017; Care *et al*., 2019; Leinonen *et al*., 2020; Shen *et al*., 2020). For example, Care *et al*. ablated 50% of rods in adult mice and uncovered functional compensation at the level of retinal interneurons, which recovered retinal output to near-normal levels (Care *et al*., 2020). Contextually similar conclusions of homeostatic adaptation to a partial retinal injury were drawn from rod/cone, cone, or rod bipolar cell ablation studies in adult rabbit, ground squirrel, and mouse retinas (Johnson *et al*., 2017; Beier *et al*., 2018; Care *et al*., 2019; Shen *et al*., 2020). We showed in the *Rho*^P23H/WT^ mouse, representing a common form of human autosomal dominant RP, increased sensitivity to visual inputs at the photoreceptor-to-bipolar cell synapse and maintained rod-driven contrast sensitivity regardless of rod population loss of more than 50% (Leinonen *et al*., 2020). Importantly, functional adaptation to retinal stress has also been observed in glaucoma models, wherein increased excitability of RGCs and dLGN neurons may mitigate the loss of axon function caused by elevated intraocular pressure (Calkins, 2021; Van Hook *et al*., 2021).

Altogether, accumulating evidence suggests that retinal remodeling upon photoreceptor degeneration may not solely possess disruptive elements, but may initially help the visual system by adapting to sensory function loss. As these countering elements seem to temporally overlap, it will be of utmost importance to uncover what are the original molecular triggers for the retinal rewiring and if these signals divide into more than one downstream pathway. Will it be possible to pharmacologically intervene to promote neural adaptation while simultaneously suppressing the disruptive hyperexcitability? With respect to vision restoration trials in adult humans, neural plasticity will be crucial.

## ABBREVIATIONS

AMSA: amplitude spectrum area
C50: contrast at the 50% of the maximum response
dLGN: dorsal lateral geniculate nucleus
ERG: electroretinogram
HWHH: half-width at half height
OMR: optomotor response
OPs: oscillatory potentials
RGC: retinal ganglion cell
*Rho*: rhodopsin
RP: Retinitis Pigmentosa
RD: retinal degeneration
SF: spatial frequency
TF: temporal frequency
VEP: visually evoked potential
V1: primary visual cortex
WT: wild type.

## ACKNOWLEDGMENTS

We acknowledge unrestricted grants from Research to Prevent Blindness to the Departments of Ophthalmology at UCI. We thank Dr. Vladimir Kefalov for valuable comments during manuscript preparation. We also thank our colleagues at UCI Center for Translational Vision Research and Gavin Herbert Eye Institute for helpful comments regarding this study. Dr. Palczewski is the Donald Bren Professor and the Irving H. Leopold Chair of Ophthalmology.

---

**Supplementary Figure S1.**
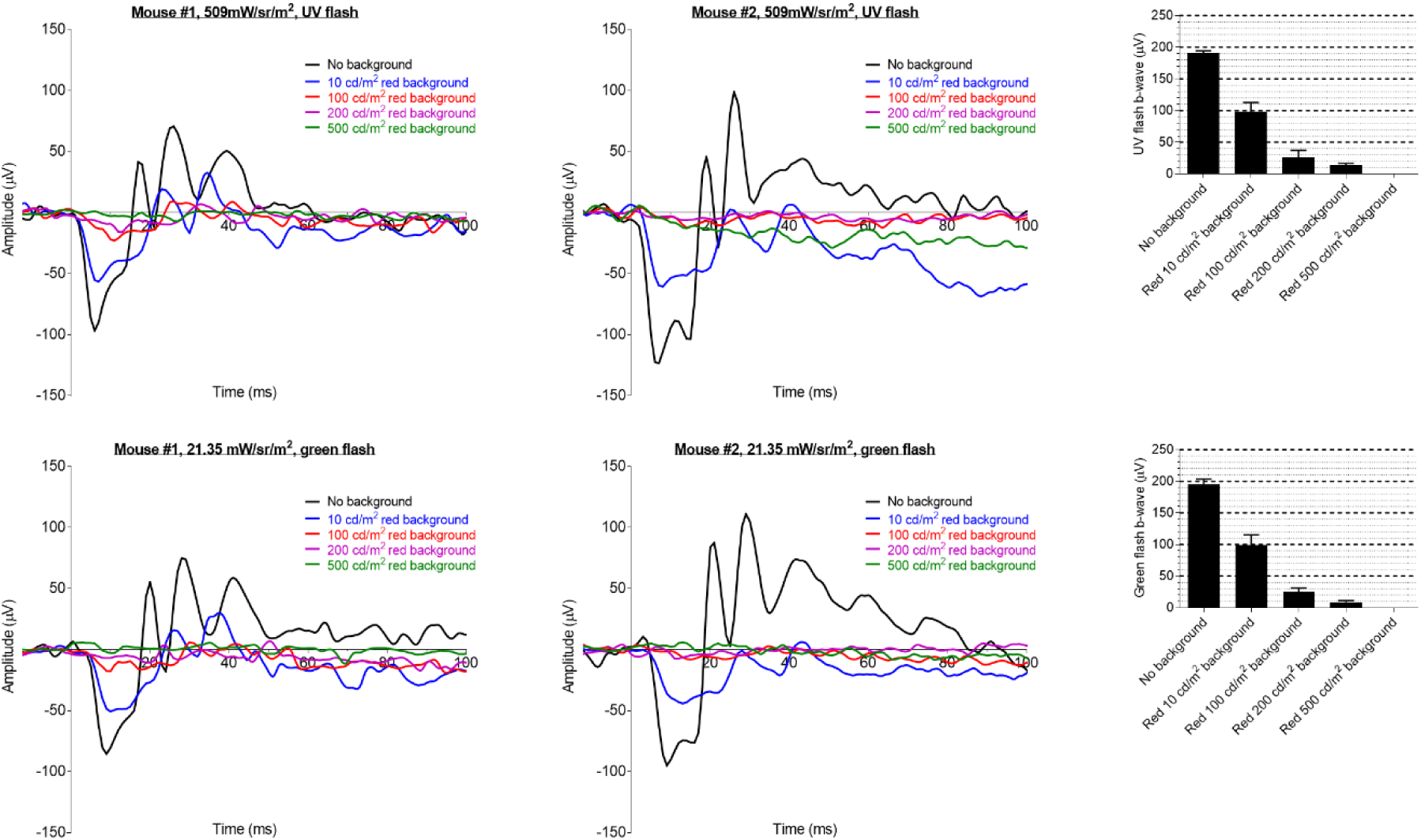
Steady red illumination suppresses rod-driven ERG in cone-transducin knockout (Gnat2^-/-^) mice. *Gnat2*^*-/-*^ mice were dark-adapted overnight before the recording. Mice were first recorded in a dark-adapted state (no background), and then steady red light at 10, 100, 200, and 500 cd/m^2^ was introduced, and ERGs were repeated in an increasing ” red light-adapted” state. Each step had an adaptation period of 60 sec. Two stimuli were presented for each step using an interstimulus interval of 90 sec. Note that the ERG response in 200 cd/m^2^ background illumination is less than 10% from the dark-adapted state, and the response in 500 cd/m^2^ background illumination cannot be distinguished from the noise.

**Supplementary Figure 2.**
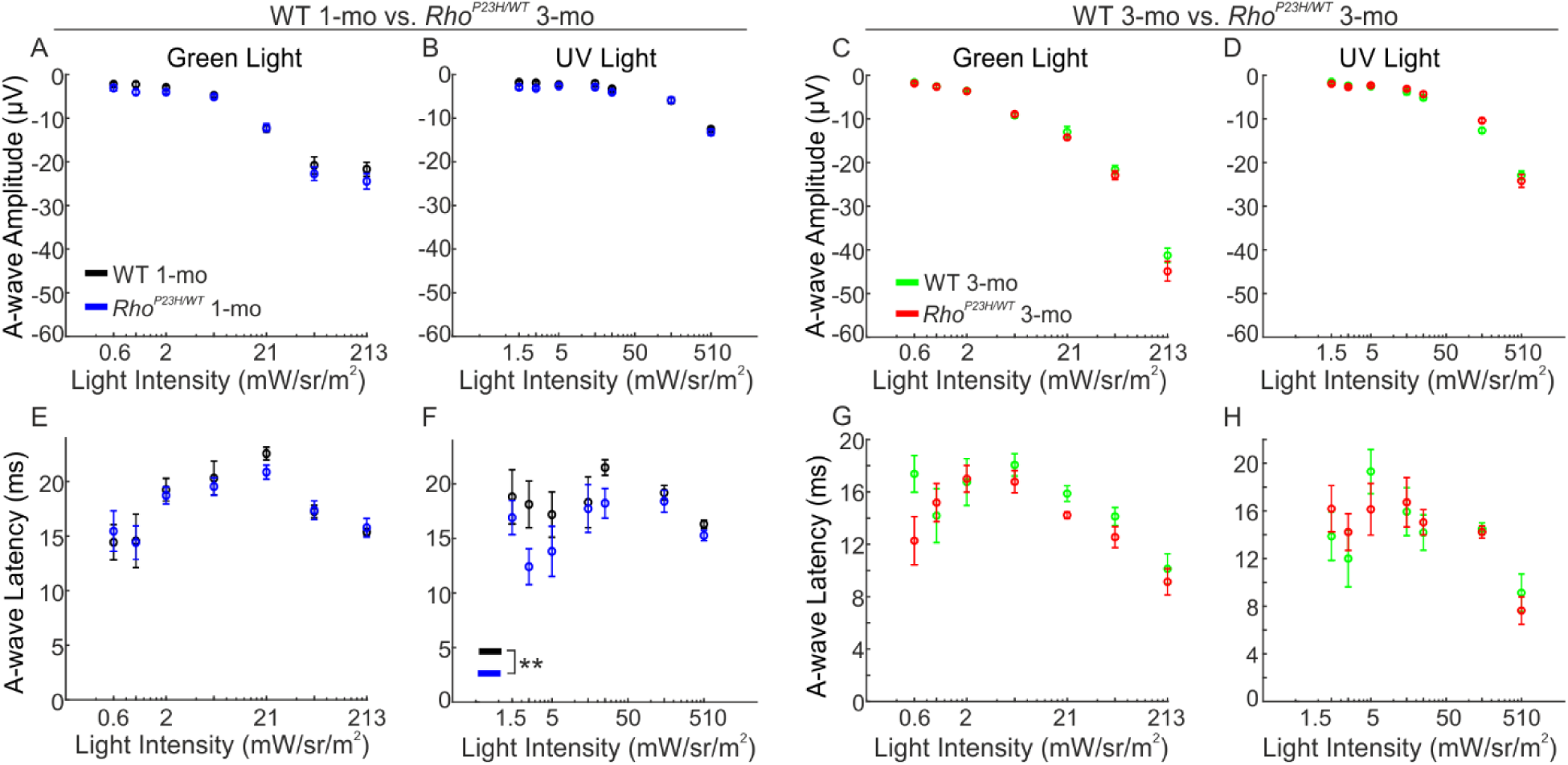
Photopic ERG a-wave amplitude and latency graphs. Panels A,B,E,F represent data from 1-month-old *Rho*^*P23H/WT*^ mice (blue, n=11) and WT mice (black, n=8). Panels C,D,G,H represent data from 3-month-old *Rho*^*P23H/WT*^ mice (red, n=11) and WT mice (green, n=11). The RM ANOVA between-subjects effect revealed that the a-wave peak occurred significantly faster in 1-month-old *Rho*^*P23H/WT*^ mice compared to WT (graph F; RM ANOVA between-subjects effect: *P*<0.01). There were no statistically significant differences for any of the other conditions as measured by RM ANOVA.

**Supplementary Figure 3.**
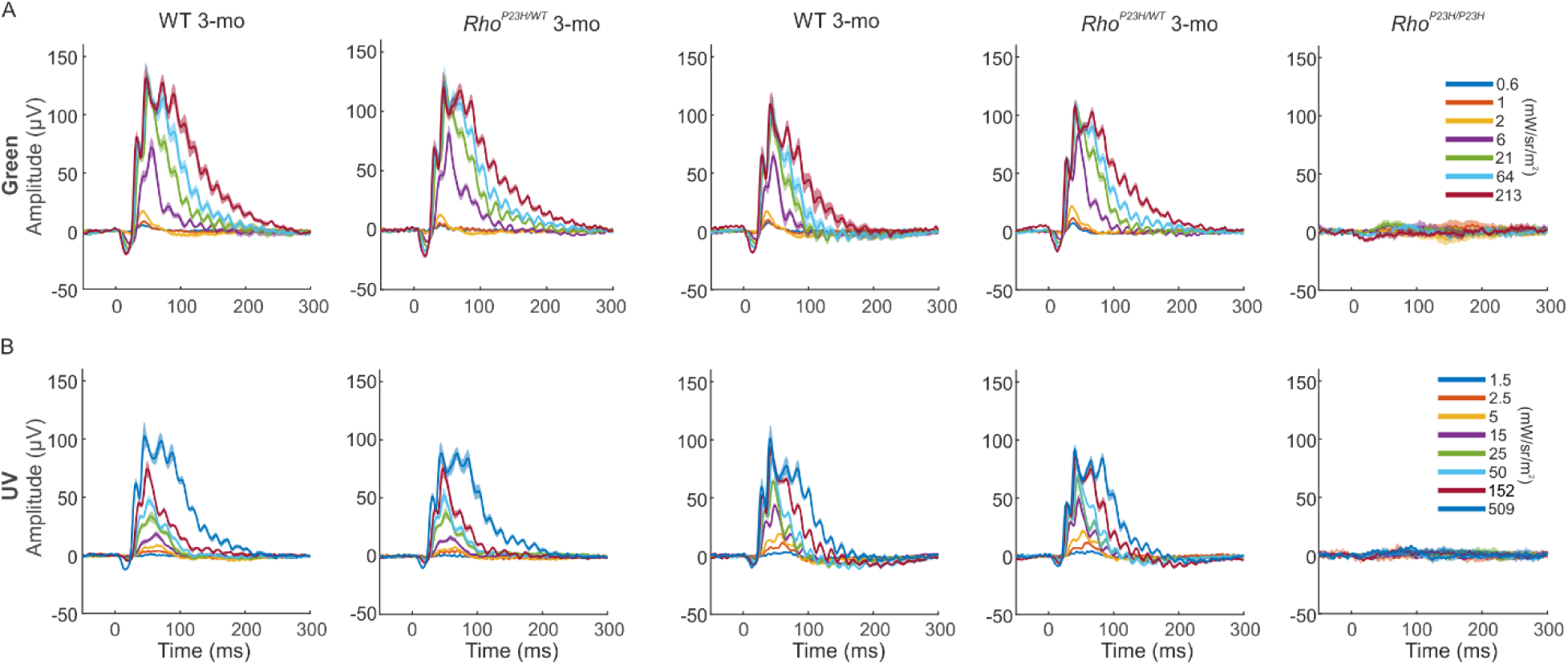
Group-averaged ERG waveforms for every stimulus condition for each animal group. (A) Green flash responses. (B) UV flash responses.

**Supplementary Figure 4.**
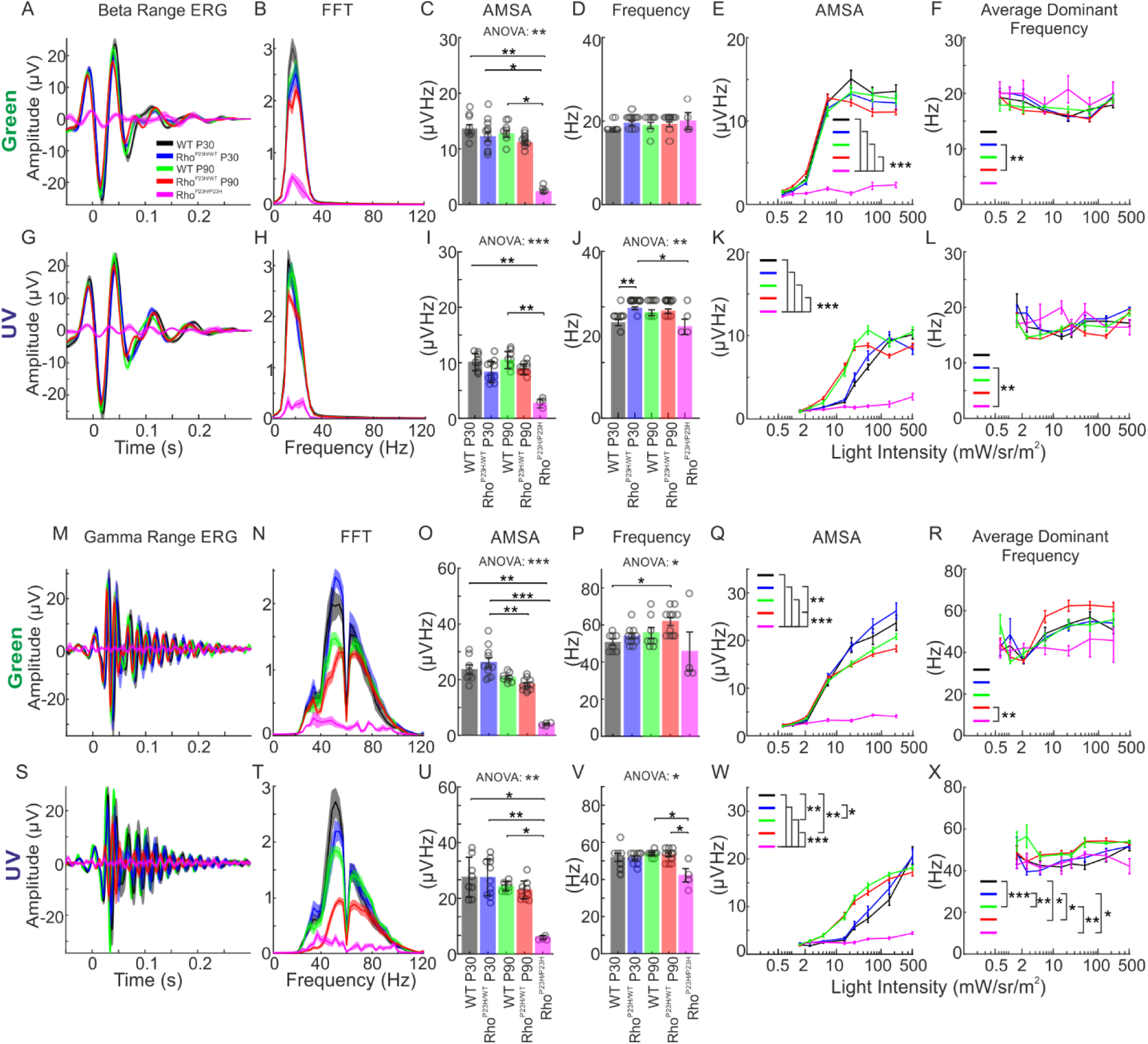
Spectral dissection of oscillatory potentials from the ERG responses. (A) Filtered ERG response at beta range in response to 214 mW·sr/m^2^ green flash. (B) Amplitude spectra corresponding to signals from A. (C) The amplitude spectrum area (AMSA) calculated from B. (D) The dominant frequency from response in A. (E) AMSA at all green flash intensities. (F) Dominant response frequencies at all green flash intensities (G) Filtered ERG response at beta range in response to 509 mW·sr/m^2^ UV flash. (H) Amplitude spectra corresponding to signals from G. (I) The AMSA calculated from H. (J) The dominant frequency from the response in G. (K) AMSA at all UV flash intensities. (L) Dominant response frequencies at all UV flash intensities. (M) Filtered ERG response at gamma range in response to 214 mW·sr/m^2^ green flash. (N) Amplitude spectra corresponding to signals from M. (O) The AMSA calculated from N. (P) The dominant frequency from response in M. (Q) AMSA at all green flash intensities. (R) Dominant response frequencies at all green flash intensities (S) Filtered ERG response at gamma range in response to 509 mW·sr/m^2^ UV flash. (T) Amplitude spectra corresponding to signals from S. (U) The AMSA calculated from T. (V) The dominant frequency from the response in S. (W) AMSA at all UV flash intensities. (X) Dominant response frequencies at all UV flash intensities. Statistical analysis was performed by the Kruskal-Wallis test followed by Dunn-Sidak post hoc tests: \**P*<0.05, \*\**P*<0.01, \*\*\**P*<0.001.

**Supplementary Table 1.** Conversion of luminous energy to radiance units. Conversion coefficients are as follows: 0.002135 W/sr/m^2^ = 1 photopic cd·s/m^2^ for the green light; and 0.050915 W/sr/m^2^ = 1 photopic cd·s/m^2^ for the UV light. <u>Note</u>: The Diagnosys Espion ERG software uses luminance units by default.

| UV,<br>cd·s/m<sup>2</sup> | UV,<br>W/sr/m<sup>2</sup> | UV,<br>mW/sr/m<sup>2</sup> | Green,<br>cd·s/m<sup>2</sup> | Green,<br>W/sr/m<sup>2</sup> | Green,<br>mW/sr/m<sup>2</sup> |
| --- | --- | --- | --- | --- | --- |
| 0.03 | 0.00152748 | **1.53** | 0.3 | 0.000640553 | **0.64** |
| 0.05 | 0.0025458 | **2.55** | 0.5 | 0.001067589 | **1.07** |
| 0.1 | 0.00509159 | **5.09** | 1 | 0.002135178 | **2.14** |
| 0.3 | 0.01527478 | **15.27** | 3 | 0.006405533 | **6.41** |
| 0.5 | 0.02545796 | **25.46** | 10 | 0.021351776 | **21.35** |
| 1 | 0.05091592 | **50.92** | 30 | 0.064055328 | **64.06** |
| 3 | 0.15274777 | **152.75** | 100 | 0.21351776 | **213.52** |
| 10 | 0.50915923 | **509.16** |  |  |  |

